# A novel genetic mouse model of fatal aortic dissection reveals massive inflammatory cell infiltration in the thoracic aorta

**DOI:** 10.1101/2024.05.02.592287

**Authors:** Kenichi Kimura, Eri Motoyama, Sachiko Kanki, Keiichi Asano, Md Al Amin Sheikh, Maria Thea Rane Dela Cruz Clarin, Erna Raja, Mariko Takeda, Ryutaro Ishii, Kazuya Murata, Violette Deleeuw, Patrick Sips, Laura Muiño Mosquera, Julie De Backer, Seiya Mizuno, Lynn Y Sakai, Tomoyuki Nakamura, Hiromi Yanagisawa

## Abstract

**Background:** Aortic dissection (AD) is the separation of medial layers of the aorta and is a major cause of death in patients with connective tissue disorders such as Marfan syndrome. However, molecular triggers instigating AD, its temporospatial progression, and how vascular cells in each vessel layer interact and participate in the pathological process remain incompletely understood. To unravel the underlying molecular mechanism of AD, we generated a spontaneous AD mouse model.

**Methods:** We incorporated a novel missense variant (p.G234D) in *FBN1*, the gene for fibrillin-1, identified in a non-syndromic familial AD patient into mice using CRISPR/Cas9 system. We performed histopathological analyses of the aortic lesions by histology, immunofluorescence staining, electron microscopy, synchrotron-based imaging and single-cell (sc)RNA-sequencing. Biochemical analysis was performed to examine the binding capacity of mutant human FBN1G234D protein to latent Tgfβ binding proteins (LTBPs), and signaling pathways in the mutant aortic wall were examined by western blot analysis.

**Results:** 50% of the *Fbn1*^G234D/G234D^ mutant mice died within 5 weeks of age from multiple intimomedial tears that expanded longitudinally and progressed to aortic rupture accompanied by massive immune cell infiltration. scRNA-sequencing, validated by immunostaining, revealed a significant increase in MHC class II-positive pro-inflammatory macrophages and monocytes at the site of intima tears with upregulation of MMP2/9 and marked disruption of elastic lamina. Subendothelial matrices, such as type IV collagen and laminin, expanded into the medial layer, where fibronectin expression was highly upregulated. *Fbn1*^G234D/G234D^ endothelial cells exhibited altered mechanosensing with loss of parallel alignment to blood flow and upregulation of VCAM-1 and ICAM-1, all of which likely contributed to the infiltration of immune cells. Biochemically, FBN1G234D lost the ability to bind to latent TGFβ binding protein (LTBP)-1, -2, and -4, resulting in the downregulation of TGFβ signaling in the aortic wall.

**Conclusions:** We show that dynamic interactions involving endothelial cells (ECs) and macrophages/monocytes in the intima, where the ECM microenvironment is altered with the reduced TGFβ signaling, contributes to the initiation of AD. Our novel AD mouse model provides a unique opportunity to identify target molecules involved in the intimomedial tears that can be utilized for development of therapeutic strategies.

## Introduction

Aortic dissection (AD), tears that separate the medial layers of the aorta, can rapidly progress to aortic rupture or cardiac failure, leading to high mortality without timely surgical treatment ^1^. Since the risk of AD often increases in thoracic aortic aneurysm (TAA) with progressive weakening of the aortic wall, it can be regarded as the end-stage consequence of AA ^2^. However, AD can also occur without aneurysms or in mildly dilated aortas, and it accounts for early mortality among patients with connective tissue disorders ^3,4^. Therefore, it is crucial to investigate the etiology of AD separately from that of AA in order to develop therapeutic strategies tailored to target the initiation and/or progression of AD. AD has been extensively studied by analyzing the pathology of human surgical aortic tissue samples, and intimal tear or laceration was observed in most cases ^5^. However, the temporospatial progression of AD and how vascular cells in each vessel layer interact and participate in the pathological process have not been fully understood.

Currently available AD animal models are largely divided into three types: First, chemically-induced AD models using beta amino propionitrile (BAPN), an irreversible lysyl oxidase inhibitor that inhibits crosslinking of collagen and elastin, with or without angiotensin II (AngII) in normolipidemia or hyperlipidemia condition ^6^. The chemical induction causes rapid destruction of the vessel wall and induces dissection. Second, genetic thoracic aortic aneurysm and dissection (TAAD) models with mutations in the fibrillin-1 (FBN1) gene, including the *Fbn1*^mgR/mgR^ mouse which contains hypomorphic alleles of *Fbn1* that develop AA followed by rupture or AD, and the *Fbn1*^GT-8/H1Δ^ mouse which contains a truncated allele of *Fbn1* and an allele lacking the first hybrid domain of fibrillin-1 that mediates the binding to latent TGFβ-binding proteins (LTBPs) ^7^. Third, vascular Ehlers-Danlos syndrome mouse models carrying a knock-in allele for *Col3a1* point mutations (*Col3a1*^G209S/+^, *Col3a1*^G938D/+^) that develop AD without aneurysms or inflammation ^8^ and a *Col3a1* knockout mouse ^9^. It is currently unknown whether there is a common pathological process that leads to dissection in these different types of models besides the underlying weakening of the vessel wall.

Fibrillin-1 is a major component of microfibrils that serve as a scaffold for elastic fiber assembly and provides a hub for elastic fiber-associated proteins such as fibulin-4 (FBLN4), fibulin-5 (FBLN5), LTBPs, and fibronectin, all of which are essential for the formation of elastic fibers (reviewed in ^10^). Fibrillin-1 also regulates intercellular signaling mediated by transforming growth factor beta (TGFβ) and bone morphogenetic proteins (BMP)s through binding of the large latent TGFβ complex or by directly binding to the prodomains of BMPs. The contribution of fibrillin-1 to the regulation of bioavailability of growth factors has offered different interpretations using different mouse models. Increased TGFβ signaling was found in *Fbn1* mutant mice (*Fbn1*^mgΔ/mgΔ^, *Fbn1*^C1041G/+^), and the rescue of aortic and lung phenotypes by TGFβ neutralizing antibodies suggested that TGFβ was the culprit for the destruction of the aortic wall ^11,12^. Our recent studies using heterozygous *Fbn1* mutant mice carrying the deletion of the LTBP-1 binding domain (*Fbn1*^H1Δ/+^) show microdissection of the elastic lamina in the thoracic ascending aorta with normal aortic diameters and reduced TGFβ signaling, whereas in *Fbn1*^GT-8^/^H1Δ^ mice, TGFβ-mediated signaling is significantly increased and AA progresses rapidly to rupture ^7^. In *Fbn1*^H1Δ/+^ mice, mast cells were identified in the adventitia of the thoracic ascending aorta with a reduced amount of TGFβ-downstream signals, suggesting an inverse relationship between the level of TGFβ signaling and inflammation in the aorta.

It has been reported that macrophages constitute the predominant cell type in aortopathy (Reviewed by ^13,14^). Macrophages are categorized into two primary phenotypes– the classically activated M1 inflammation-inducing macrophages and the alternatively activated M2 anti-inflammatory macrophages. When activated, pro-inflammatory macrophages secrete inflammatory cytokines such as tumor necrosis factor (TNF)-α, interleukin (IL)-1β, IL-6, monocyte chemotactic protein (MCP1), and upregulate inducible nitric oxide synthase (iNOS). Conversely, substances generated by anti-inflammatory macrophages such as IL-10 and TGFβ and reduced expression of MHC class II resolve inflammation and contribute to the repair process ^15^.

In this study, we report a novel AD mouse model carrying a likely pathogenic *FBN1* variant identified in non-syndromic familial AD patients and investigate the molecular mechanism of initial pathogenic events in the aortic wall. We show that dynamic interactions involving endothelial cells (ECs) and macrophages/monocytes in the intima where the altered ECM microenvironment with the reduced TGFβ signaling contribute to the development of AD.

## Methods

Detailed descriptions are provided in the Online Data Supplement.

### Mice

*Fbn1*^G234D^ mutant mice were developed by using the CRISPR/Cas9 genome editing system. A targeting site within the exon 7 of mouse *Fbn1* was selected using the CRISPRdirect web server (http://crispr.dbcls.jp/).

## Results

### Generation of a new mouse model based on a pathogenic variant identified in human familial aortic dissection

A male patient in his 30’s suffered from Stanford type A AD without root or ascending aortic aneurysm. Four affected family members in two generations also had isolated AD (Figure 1A). The proband was found to be heterozygous for the missense variant (c.701 G>A, p.234 Gly>Asp) in the first hybrid domain of fibrillin-1. Amino acid 234 is a glycine residue that is conserved in all hybrid domains, in the 8-cysteine domain and in the corresponding region of EGF-like domains. However, he lacked typical Marfanoid features such as lens dislocation or systemic features (Figure 1B). To investigate the causal association between the missense variant and AD, we developed a mouse strain carrying the corresponding variant using the CRISPR/Cas9 system. Seventeen of 59 surviving founders had mutant alleles, of which two lines were used for further analysis. Subsequent intercrosses followed a normal Mendelian inheritance (Figure S1A). *Fbn1*^G234D*/+*^ heterozygous mutant mice were healthy and had a full life span without aortic defects (Figure S1B and S1C). In contrast, *Fbn1*^G234D/G234D^ homozygous mutant mice died suddenly starting from 3 weeks of age, with 50% mortality by 5 weeks of age (Figure 1C). The survival rate showed no sexual dimorphism and no difference in body weight was observed between female wild-type (WT) and mutant mice; however, male mutant mice had slightly lower body weight compared to WT mice at 5 weeks of age (Figure S1D). Tortuosity of the aorta was observed in mutant mice with 100% penetrance at 3 weeks (n=56) and kyphosis appeared in a few mutant mice at 5 weeks of age (4/35). Tight skin was noted at 7 days of age in all the mutant (n=13).

**Figure 1.**
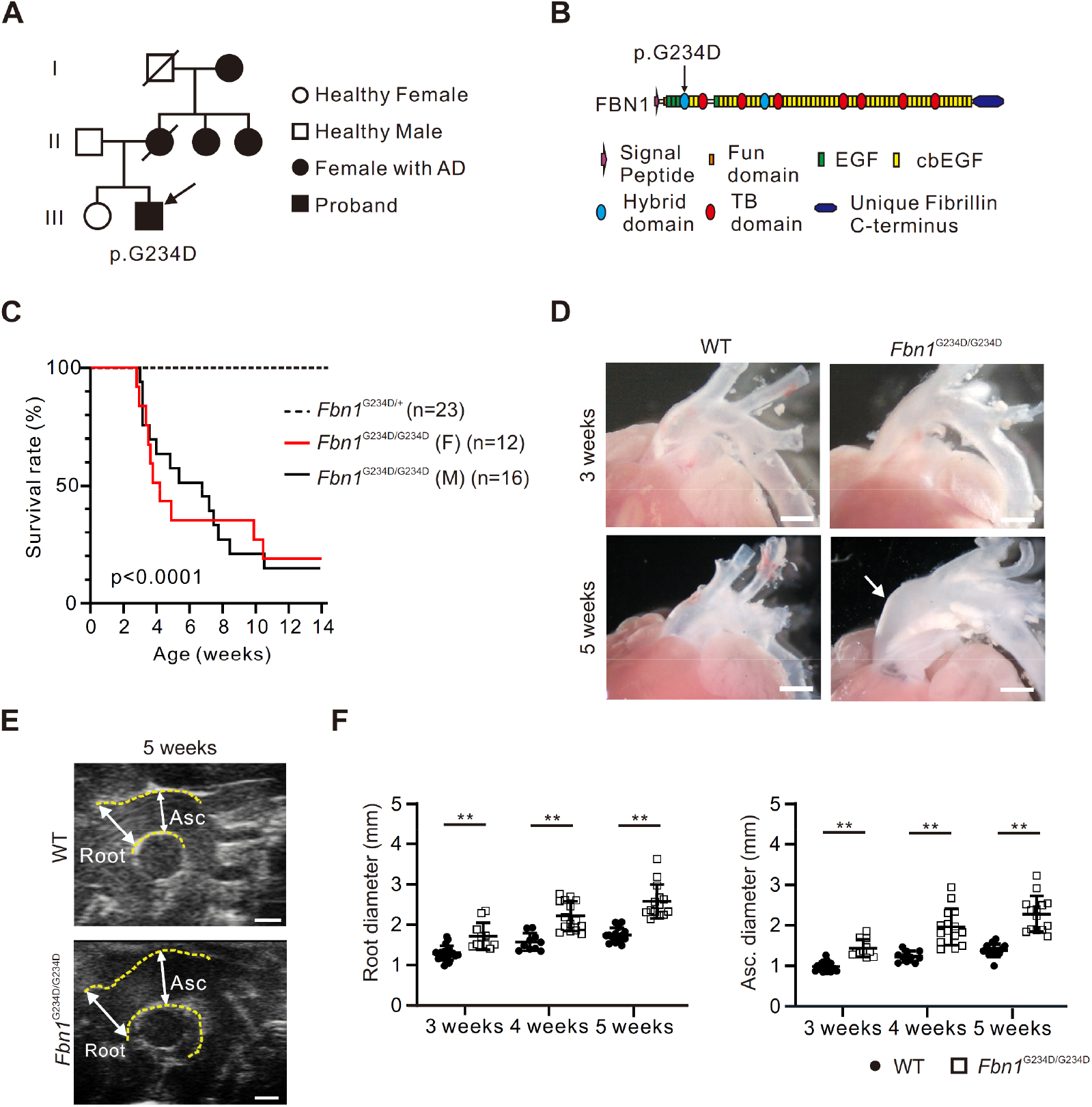
Familial aortic dissection with a novel *FBN1* missense variant. (A) Family tree of the young male patient with a novel *FBN1* missense variant. (B) Schematic image of fibrillin-1 (FBN1) domains. The missense variant (c.701 G>A, p.234 Gly>Asp, arrow) in the first hybrid domain was identified in the *FBN1* gene. (C) Kaplan-Meier survival curve of *Fbn1* mutant mice (n=23), *Fbn1*^G234D/G234D^ female (F) mice (n=12), *Fbn1*^G234D/G234D^ male (M) mice (n=16). Heterozygous v.s. homozygous mutant mice, p<0.0001, Log-rank test. (D) Gross images of WT and *Fbn1*^G234D/G234D^ aortas at 3 and 5 weeks of age. Note the dilated ascending aorta in 5-week-old mutant mouse (arrow). (E) Representative images of echocardiography of aortas from WT and mutant mice at 5 weeks of age. (F) Quantification of echocardiography showing a significant increase in maximum diastolic diameter in root and ascending aortas. Mean ± SEM. 3 weeks WT: n=19, *Fbn1*^G234D/G234D^: n=11; 4 weeks WT: n=10, *Fbn1*^G234D/G234D^: n=15; 5 weeks WT: n=14, *Fbn1*^G234D/G234D^: n=14. **p<0.01, one-way ANOVA. Bars are 1 mm (D and E).

To examine the cause of death in mutant mice, we performed macroscopic *ex vivo* observations at 3 and 5 weeks of age (Figure 1D). At 3 weeks, there were no discernable differences between WT and mutant mice, but dilatation of the ascending aorta was observed in mutant mice at 5 weeks of age (arrow in Figure 1D). *In vivo* cardiovascular ultrasound of the thoracic aorta confirmed increased diameters of the aortic root and ascending aorta starting at 3 weeks of age (Figure 1E-F). Upon necropsy, intrapleural hemorrhage was observed with massive blood attached to the ascending aorta and aortic arch, but not in the abdominal aorta, indicating thoracic AD and rupture (Figure S1E). Study of ventricular function showed a normal ejection fraction (EF), whereas the transmitral doppler flow velocity pattern showed significantly reduced early diastolic to atrial systolic flow velocity (E/A) in the mutant mice at 5 weeks of age (Figure S1F and S1G).

### Spontaneous aortic dissection with intimomedial tears in *Fbn1*^G234D/G234D^ mice

To investigate pathological processes in *Fbn1*^G234D/G234D^ aortas, we examined serial cross-sections of the aortas stained with HE for routine histology and Hart’s for elastic fibers. At 3 weeks, the aortic wall was largely intact in *Fbn1*^G234D/G234D^ mice, and the wall thickness was comparable between WT and mutant mice (Figures 2A-2C). However, the elastic laminae in the mutant ascending aortas contained breaks, which were already evident at 1 week old (Figure 2D). The ultrastructural analysis revealed that elastic laminae of the WT ascending aortas were continuous and elastin-contractile units were being formed, providing a direct connection between elastic fibers and smooth muscle cells (SMCs, arrows in Figure 2E). In contrast, elastic laminae of *Fbn1*^G234D/G234D^ aortas were disrupted with loss of contacts with SMCs (arrowheads in Figure 2E) and the orientation of SMCs was disorganized, suggesting the underlying developmental defects of elastic fibers due to the mutation in fibrillin-1. The fragmentation of elastic laminae became more prominent as mutant mice matured. Some immune cells were detected in the adventitia of the *Fbn1*^G234D/G234D^ aortas (arrowheads in Figure 2B). At 5 weeks old, multiple intimomedial tears were found in the mutant aortas (arrowheads in Figure 2F), ranging from breaks in the internal elastic lamina (IEL) to nearly complete breaks throughout the medial layers with the presence of thin external elastic lamina (Figure 2G). There was a disruption in the continuity of medial layers and massive infiltrates were observed in the adventitia (arrows in Figure 2G), and in some cases, medial separation was clearly observed (arrowhead in Figure S1H).

**Figure 2.**
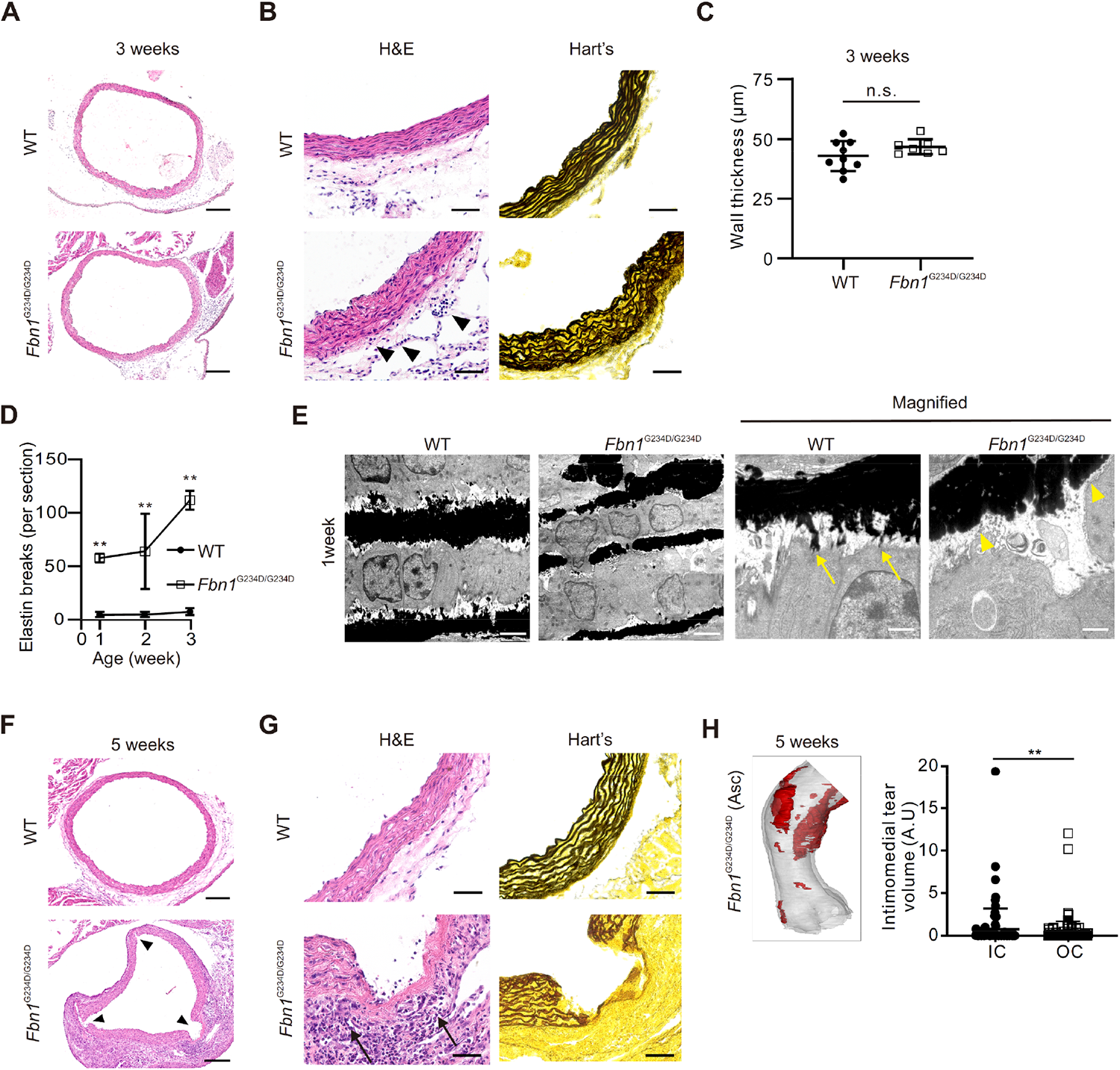
Aortic dissection with intimomedial tears in the ascending aortas of Fbn1^G234D/G234D^ mouse. (A, B) Histological images of WT and *Fbn1*^G234D/G234D^ aortas with H&E and Hart’s staining at 3 weeks of age. Elastin breaks and accumulation of immune cells (arrowheads) are observed in the aorta from *Fbn1*^G234D/G234D^ mice. (C) The morphometric analysis of wall thickness displays no significant differences between WT and *Fbn1*^G234D/G234D^ aortas. WT: n=9, *Fbn1*^G234D/G234D^: n=7, n.s.: not significant, unpaired *t*-test. (D) The quantification of elastin breaks in the aortas from WT and *Fbn1*^G234D/G234D^ mice at 1 to 3 weeks of age. Mean ± SEM. At 1 week WT: n=3, *Fbn1*^G234D/G234D^: n=3; 2 weeks WT: n=6, *Fbn1*^G234D/G234D^: n=5; 3 weeks WT: n=9, *Fbn1*^G234D/G234D^: n=7. **p<0.01, one-way ANOVA. (E) Electron microscopic images of ascending aorta obtained from WT and *Fbn1*^G234D/G234D^ mice at 1 week of age. In WT aorta, continuous elastic laminae were being formed with the elastin-contractile unit (arrows), whereas elastic laminae of *Fbn1*^G234D/G234D^ aortas were disrupted with loss of contacts with smooth muscle cells (arrowheads). (F, G) Histological images of WT and *Fbn1*^G234D/G234D^ aortas with H&E and Hart’s staining at 5 weeks of age. There were multiple intimomedial tears (arrowheads) in the aorta from *Fbn1*^G234D/G234D^ mice (F). The continuity of medial layers was disrupted and red blood cells were detected within the medial layers (arrows, G). (H) 3D structure of intimomedial tears and quantification of its volume normalized by ascending volume in the aorta from *Fbn1*^G234D/G234D^ mice at 5 weeks. Multiple lesions (red) with disrupted medial layers expanded longitudinally. IC: inner curvature, OC: outer curvature. Disrupted lesions (n=236) from 10 mice were analyzed. **p<0.01, Mann-Whitney U test. Bars are 200 µm (A and F), 50 µm (B and G), 2 µm (E) and 1 µm (magnified in E).

To examine the spatial aspect of intimomidial tears and to search for preferential locations, i.e., inner curvature (IC) vs. outer curvature (OC), we performed synchrotron-based phase contrast X-ray tomographic microscopy imaging analysis using 3-and 5-week-old ascending thoracic aortas. This high-resolution analysis allows us to visualize aortic lesions in both transverse and longitudinal views with single-lamina resolution without serial sectioning, enabling 3-dimensional (3D) segmentation ^16–18^ (Figure S2A). At 3 weeks old, *Fbn1*^G234D/G234D^ aorta was slightly dilated, consistent with echocardiography data, and some exhibited small intimal tears (Figure S2B). At 5 weeks old, multiple lesions with disrupted medial layers expanded along the long axis, mostly observed in the ascending aorta (Figure 2H). In addition, a significant preference toward the disruption in IC was observed. Taken together, microscopic elastic damages were already present before 3 weeks of age in *Fbn1*^G234D/G234D^ aortas, and intimomedial tears rapidly progressed to AD and aortic rupture, resulting in the death of *Fbn1*^G234D/G234D^ mice. These data support that the early aortic pathology in the mutant mice recapitulates human AD (hereafter termed AD mice).

### AD aortas exhibit compromised ECM organization

Microfibrils serve as scaffolds for elastic fiber assembly and stability. We first examined the *Fbn1* mRNA expression in primary SMCs isolated from 2-week-old WT and AD mice before massive immune cells were seen in AD aortas. No difference was observed in *Fbn1* mRNA levels between WT and AD SMCs (Figure 3A). We then assessed ECM synthesis in SMCs by elastogenesis assay, where confluent cells were cultured for 10 days and the formation of elastin-positive fibers was assessed. From day 5 to day 10, WT SMCs showed fibrous staining positive for fibronectin, fibrillin-1 and elastin, whereas much less elastin fibers were detected in AD SMCs at day 10 (Figures 3B and 3C). AD SMCs required an additional 6 days to form elastin fibers (data not shown). To examine ECM formation in vivo, immunostaining analysis was performed on the aorta. As shown in Figure 3D, a substantial decrease in fibrillin-1 immunostaining in the aortic media of AD mice was observed compared to WT mice at 3 weeks of age. This reduction was observed as early as postnatal day 3 and remained consistent at 3 weeks old (Figure 3D and S3A). The mutant fibrillin-1 fibril assembly and ECM production in SMCs were compromised possibly due to delayed secretion, impaired processes of elastogenesis, or accelerated degradation. In AD adventitia, reduction in the fibrillin-1 immunostaining was not as severe as that of media, suggesting a distinct mechanism for fibrillin-1 elastic fiber assembly between SMCs and adventitial fibroblasts. Taken together, the mutant fibrillin-1 alters ECM dynamics in the aortic wall and compromises the structural organization of the AD aorta.

**Figure 3.**
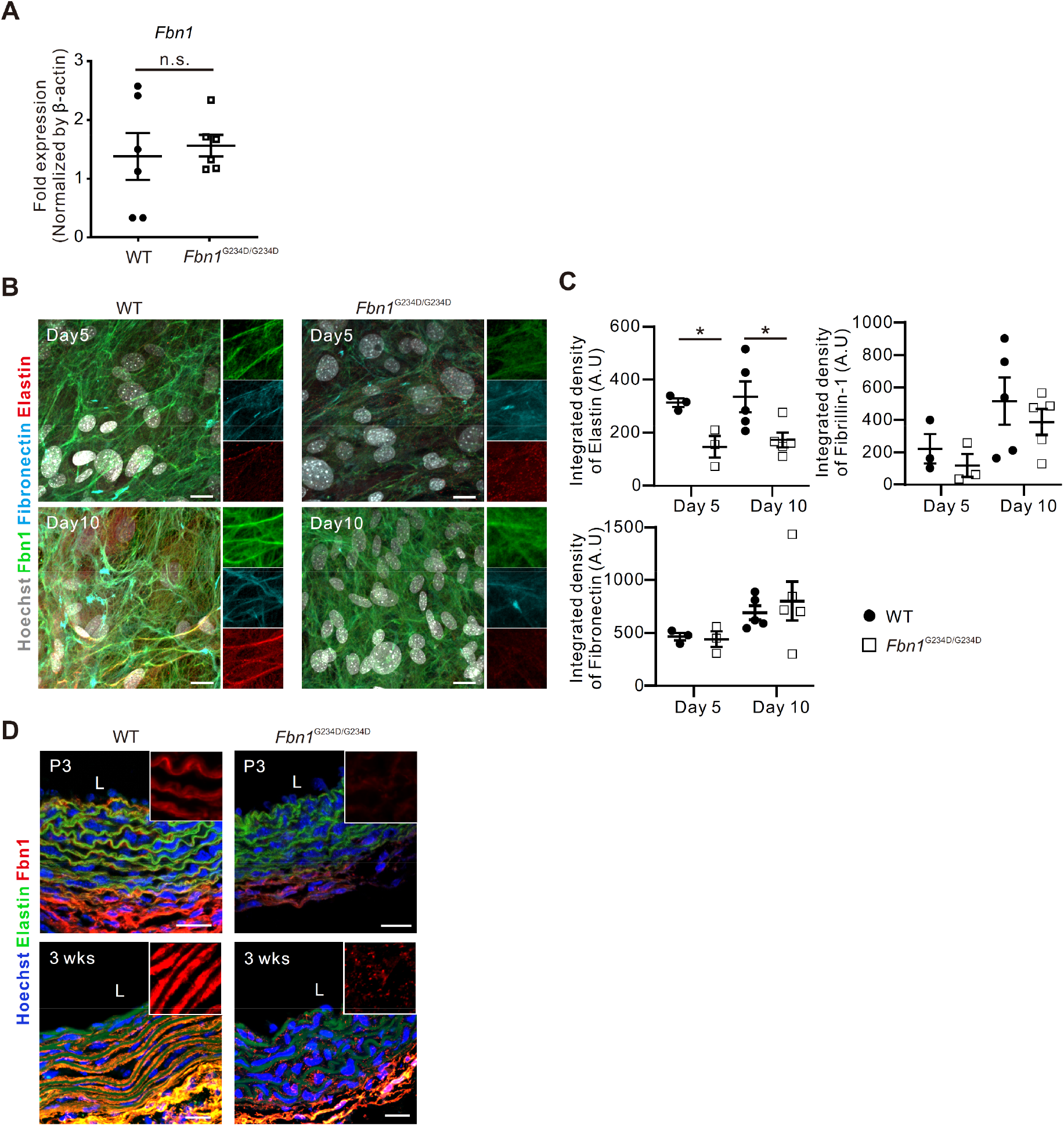
SMCs derived from *Fbn1*^G234D/G234D^ mice show defects in elastogenesis. (A) Evaluation of *Fbn1* mRNA expression by qPCR in primary SMC from 2-week-old WT and *Fbn1*^G234D/G234D^ mice. n=6 mice for each genotype. n.s.: not significant, unpaired *t*-test. (B) Representative immunofluorescence images of elastogenesis assay showing the delayed ECM formation in primary SMCs from *Fbn1*^G234D/G234D^ mice compared to WT. FN: fibronectin. n=3 (day 5), n=5 (day 10). *p<0.05, unpaired *t*-test. (C) Quantitative analysis of integrated density of elastin, fibronectin, and fibrillin-1. (D) Representative immunofluorescence images showing markedly decreased fibrillin-1 expression (red) in the media of aorta from *Fbn1*^G234D/G234D^ mice at postnatal day 3 (P3) (n=1) and 3 weeks of age (n=3). L: lumen side. Bars are 10 µm (B), 20 µm (D).

### EC abnormalities precede the accumulation of immune cells in AD aortas

To examine the relationship between the accumulation of immune cells and the progression of AD, we next investigated the dynamics of immune cells in AD ascending aortas and compared it to WT aortas by immunostaining. At 3 weeks of age, CD45-positive cells were observed in the adventitia in AD aortas (Figure 4A). At 4 weeks old, the number of CD45-positive cells increased and were observed in the intima and adventitia, but not in the medial layer (Figure 4B). The interaction between ECs and immune cells in the aorta has been well documented in association with chronic inflammation of the vascular wall such as atherosclerosis (Reviewed by ^19^). Therefore, we assessed EC morphology and EC-EC junctions by en face immunofluorescence staining with VE-cadherin in the ascending aorta. At 3 weeks of age, ECs in the WT aorta were elongated and aligned with the direction of blood flow, whereas ECs in the AD aorta exhibited disarray and cell-cell junctions were disrupted (Figure 4C). Quantification analysis indicated that AD ECs showed reduced cell surface area and increased roundness of the cells (Figures 4D, 4E, and S3B). The cell shape changes were already observed in the AD aorta from 1 week of age before immune cell accumulation (Figure S3C). Detailed analyses of en face staining surprisingly showed that CD45+ cells were within the EC layer in AD aorta at 3 weeks of age (arrowheads in Figure 4C). The 3D confocal images further showed that CD45-positive immune cells were interwoven with ECs in the intimal layer, and some infiltrated into the medial layer in the AD mice at 3 weeks of age (Figure 4F and arrowheads in 4G). To further confirm the distribution of CD45-positive cells, we generated a 3D rendering of the intima by Imaris cell imaging software. CD45-positive immune cells were observed underneath ECs (Figure 4H, green). Interestingly, some immune cells already attached to ECs at 1 week of age as judged by electron microscopy (Figure 4I).

**Figure 4.**
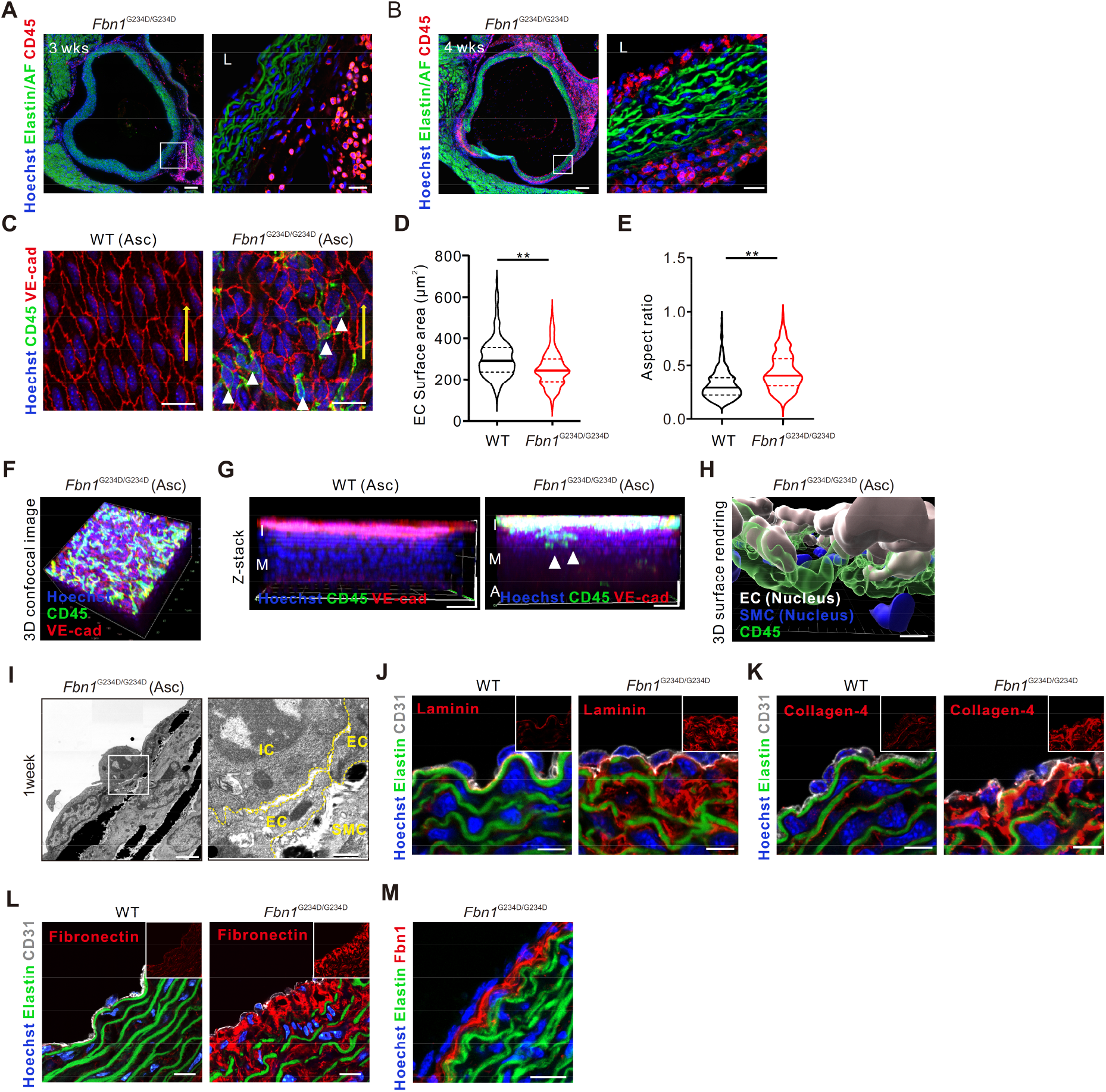
Endothelial cell (EC) abnormality in the aorta of *Fbn1* ^G234D/G234D^ mice. (A, B) Representative immunofluorescence images showing the infiltration of CD45 positive cells (red) in the intima and the adventitia of aorta from *Fbn1*^G234D/G234D^ mice at 3 weeks (n=8) (A) and 4 weeks (n=7) (B) of age. The boxed areas are shown in the magnified images to the right. (C) Representative immunofluorescence images of en face staining of the ascending aorta (Asc) show VE-cadherin positive ECs (red) fail to align with blood flow and CD45 positive cells (green, arrowheads) are observed in *Fbn1*^G234D/G234D^ mice at 3 weeks of age. (D, E) Quantitative analysis revealed that ECs from *Fbn1*^G234D/G234D^ mice exhibit reduced cell surface area (D) and increased cell roundness (E). WT: n=372 cells from 3 mice, *Fbn1*^G234D/G234D^: n=439 cells from 3 mice, **p<0.001, unpaired *t*-test. (F, G) Representative 3D confocal images show that CD45 positive cells (green) spread out in the intima (F) and infiltrate into the media (arrowheads, G) at 3 weeks of age (n=3). I: intima, M: media, A: adventitia. (H) 3D surface rendering by Imaris show immune cells distributed underneath ECs. (I) Representative electron microscopic images of the ascending aorta show attached immune cell on EC at 1 week of age. The boxed area is shown in the magnified images to the right. IC: immune cell, EC: endothelial cell, SMC: smooth muscle cell. (J, K, L, M) Representative immunofluorescence images show the abnormal expression pattern of laminin (J), collagen-4 (K), fibronectin (L), and fibrillin-1 (M) in the subendothelial layer and media at P7 (n=1) (J, K) and 5 weeks of age (n=3) (L, M). Bars are 100 µm (A, B left), 20 µm (C, G, and M), 10 µm (L), 5 µm (H, J, and K), 2 µm (I) and 500 nm (magnified in I).

To examine if subendothelial matrices affect EC morphology and/or immune cell migration to the intimal to medial layer, we examined ECMs that contribute to the microenvironment of the intimal layer. The expression of laminin and type IV collagen was markedly increased in the AD aortas and expanded into medial layers at 1 week of age (Figure 4J and 4K). As aortic lesions expanded at 5 weeks of age, fibronectin expression was observed both in the media and subendothelial space in AD aortas (Figure 4L). Furthermore, fibrillin-1 expression was upregulated in the subendothelial layer of AD aortas at 5 weeks of age (Figure 4M).

### Accumulation of inflammatory cells comprised of monocytes and macrophages

To explore cellular identities of immune cells in the aortas, we performed single-cell RNA sequencing (scRNA-seq) using isolated aortic root and ascending aorta from 5-week-old WT and AD mice. Single-cell capture and cDNA preparation were performed with the 10X Genomics Chromium platform followed by library preparation, multiplexed sequencing and analysis using the Seurat scRNA-seq analysis package in R. Using dimensionality reduction by uniform manifold approximation and projection (UMAP), we identified four major aortic cell types (SMCs, EC, fibroblasts, and immune cells) and annotated cellular identities based on canonical lineage markers (Figures S4A to S4C). In particular, the immune cell cluster was increased in the AD aortas consistent with the histological observation. To further assess the cellular identities of immune cell clusters, we performed unsupervised clustering (Figure 5A) and assigned each sub-cluster based on lineage markers (Figure 5B). There were no differences in neutrophil clusters between WT and AD mice, whereas myeloid cells (monocytes and macrophages) were markedly increased in AD aortas (Figure 5C). Within the monocytes/macrophage cell types, three distinct sub-clusters were identified (Figure 5D). The feature plot revealed the high expression of *Ccr2* and *Plac8*, both of which are highly expressed in the bone marrow and shown to be inflammatory monocytes. The second cluster contained *H2-ab1* and *H2-Dma*-positive cells compatible with inflammatory macrophages. *Mrc1* and *Lyve1* were identified in the third cluster, which are adventitial macrophages known to be protective and required for maintenance of the vessel wall ^20,21^, confirming the three distinct clustersx in the AD aortas.

**Figure 5.**
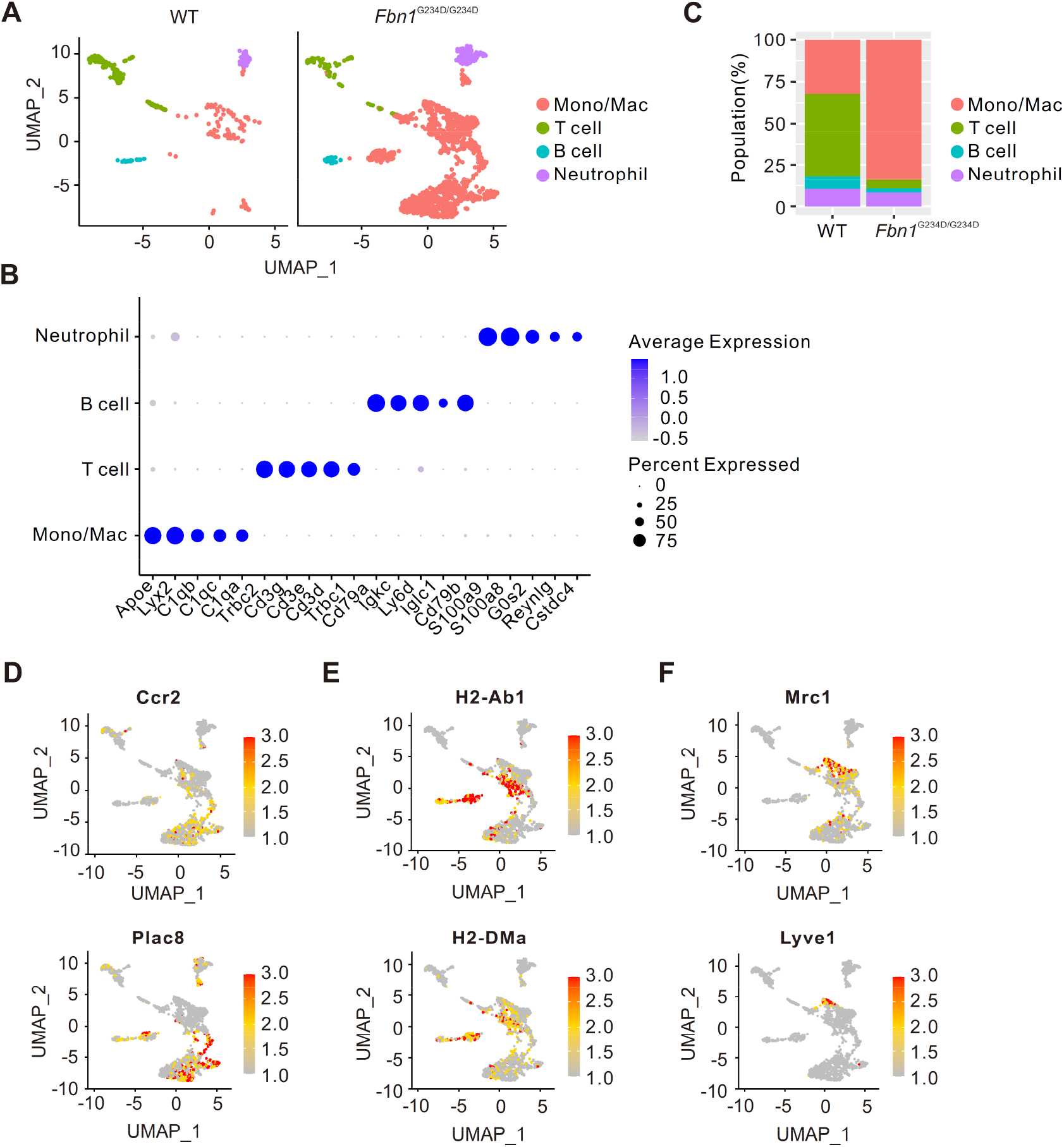
Identification of immune cell population in WT and AD assessed by scRNA-seq. (A) Uniform manifold approximation and projection (UMAP) for immune cell clusters extracted from the major cell populations. (B) The percentages of each cluster in the total number of immune cell sub-cluster. (C) A dot plot indicates the relative expression of marker genes in the distinct cell population. The dot size reflects the percentage of cells expressing the selected gene, and dot color corresponds to the level of expression. (D, E, F) Feature plots showing special distribution of monocyte (D), pro-inflammatory macrophage (E), and adventitial macrophage (F) markers in monocyte/macrophage population.

To confirm the distribution of these macrophages in the aorta, we performed immunofluorescence staining using the aortas of 5-week-old AD mice. The infiltrated immune cells in intimomedial tears were positive for CD68/MHC class II and CCR2, known to be pro-inflammatory (M1-like) macrophages, and were observed at the intima layer, whereas the cells in the adventitia were F4/80/CD206-positive repair macrophages (M2-like) (Figure 6A). In addition, in situ gelatin zymography revealed high MMP activity at the intimomedial tears in AD aortas (Figure 6B). Focal recruitment of monocytes is one of the typical cellular responses in the early stage of atherosclerosis with upregulation of adhesion molecules, including VCAM-1 and ICAM-1 on ECs ^22^. Therefore, we examined the expression of adhesion molecules in ECs in 5-week-old AD mice. In the AD ascending aorta, higher levels of ICAM-1 were observed in the non-disrupted and disrupted tunica media with intimomedial tears (Figure 6C). VCAM1 was remarkably higher in severely disrupted lesions than in mild lesions in AD aortas (Figure 6D). These observations were already seen at 1 week of age (Figures S4D and S4E).

**Figure 6.**
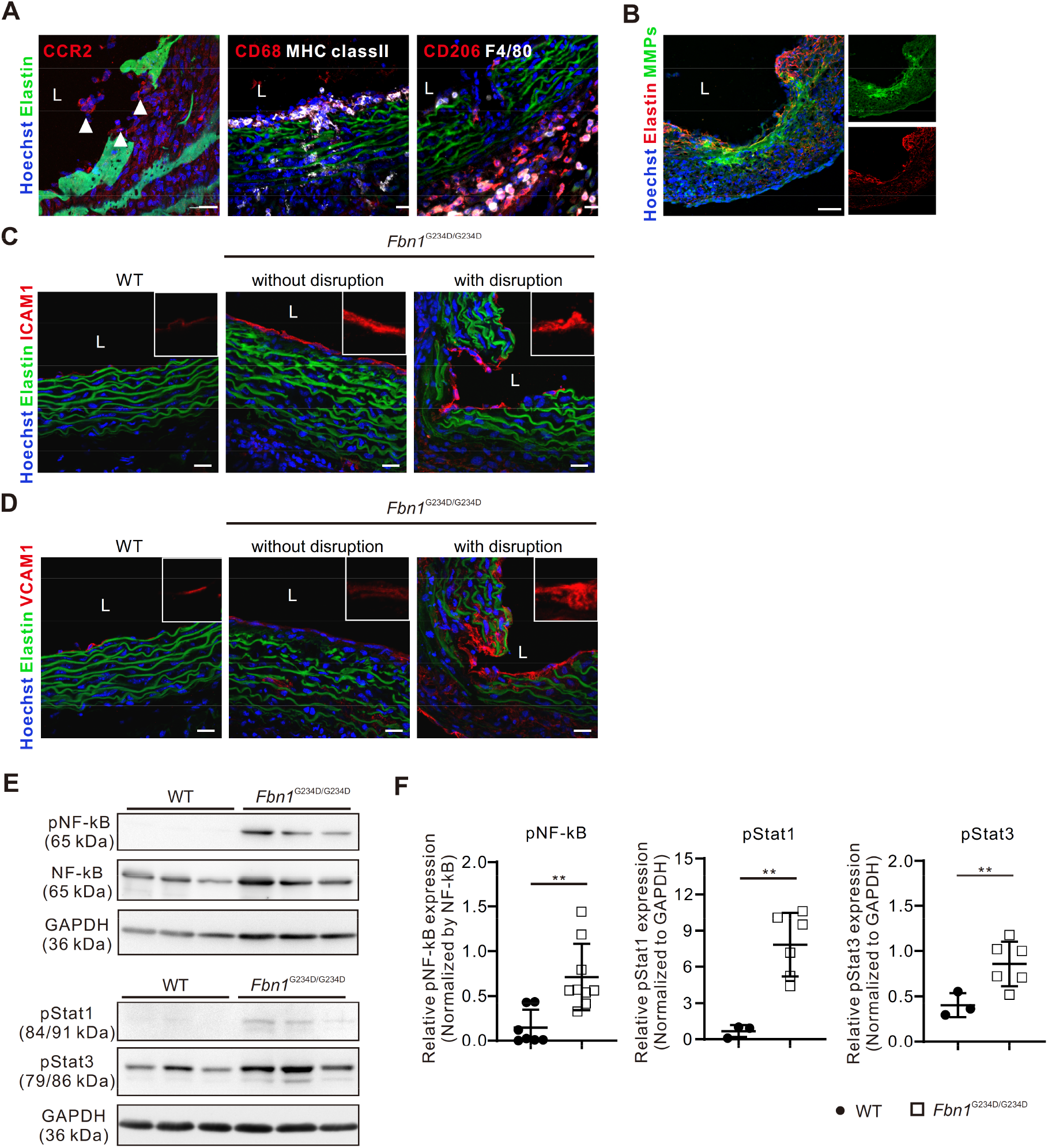
Macrophage accumulated in the aortic wall of *Fbn1* ^G234D/G234D^ mice. (A) Representative immunofluorescence images showing differential distribution of macrophages and monocyte (arrowheads) populations in the aorta from 5-week-old *Fbn1*^G234D/G234D^ mice (n=3). (B) Representative in situ zymography with DQ-gelatin showing upregulation of MMPs activity (green) in the lesions in *Fbn1*^G234D/G234D^ aortas at 5 weeks of age (n=3). (C, D) Representative immunofluorescence images showing the upregulation of ICAM-1 (C) and VCAM-1 (D) expression on the intima both with and without disruption areas at 5 weeks of age (n=3). (E, F) Western blot of total protein extracts from 3-week-old WT and *Fbn1*^G234D/G234D^ ascending aortas. Mean ± SEM, WT: n=7 (NF-κB), n= 3 (Stats), *Fbn1*^G234D/G234D^: n=9 (NF-κB), n= 6 (Stats), **p<0.01, unpaired *t* test. Bars are 20 µm (A, C, and D), 50 µm (B).

We finally investigated the underlying signaling pathway(s) responsible for inflammation in the aorta. Western blot analysis revealed that phosphorylated (p)NF-κB was significantly upregulated in the ascending aorta of 3-week-old AD mice before the onset of dissection compared to WT mice (Figures 6E and 6F). Additionally, significantly higher levels of pStat1 and pStat3 were observed in the aortas of AD mice. Taken together, these data suggested that (1) the absence of intact fibrillin-1 in the aortas has a profound effect on ECs, including morphological changes, disturbed response to blood flow, and upregulation of ICAM-1 and VCAM-1, and (2) results in the recruitment of bone marrow-derived monocyte-macrophage populations that are responsible for pro-inflammatory microenvironment in the aortic wall through the activation of NF-κB pathway. The complex interactions involving vascular cells and inflammatory cells at the intimal layer could serve as a trigger for the intimal tears and medial disruption of the AD aortas, leading to aortic dissection in AD mice.

### Glycine to aspartic acid point mutation within the first hybrid domain compromises the binding of fibrillin-1 to LTBPs and reduces canonical TGFβ pathways

The hybrid domain is known to be responsible for binding to LTBPs and fibulins ^23^, To assess the biochemical property of fibrillin-1^G234D^ protein and its influence on homeostasis of the aortic wall, we performed in vitro binding assays using purified recombinant N-terminal fibrillin-1 fragment rF23 ^24,25^ and rF23-G234D in which glycine 234 was changed to aspartic acid (Figure 7A). After individually transfecting the expression vectors into HEK293T cells, we collected the conditioned medium and cell lysates. Although rF23-G234D proteins were detected in cell lysates, the amount of protein in the conditioned medium was markedly decreased (Figure 7B), indicating that the mutation negatively impacts secretion of fibrillin-1. Using the recombinant proteins prepared from cell lysates, we performed immunoprecipitation to examine the binding of WT rF23 and rF23-G234D to LTBPs. While WT rF23 bound LTBP-1, - 2, and -4, the binding of rF23-G234D to all LTBPs was significantly impaired (Figure 7C). To exclude the possibility that cell lysates contain an inhibitory factor for protein-protein interactions, we used other known binding partners of fibrillin-1 such as fibulin-4 and fibulin-5. Surprisingly, G234D mutation did not affect binding to fibulin-4 or fibulin-5 (Figure 7D).

**Figure 7.**
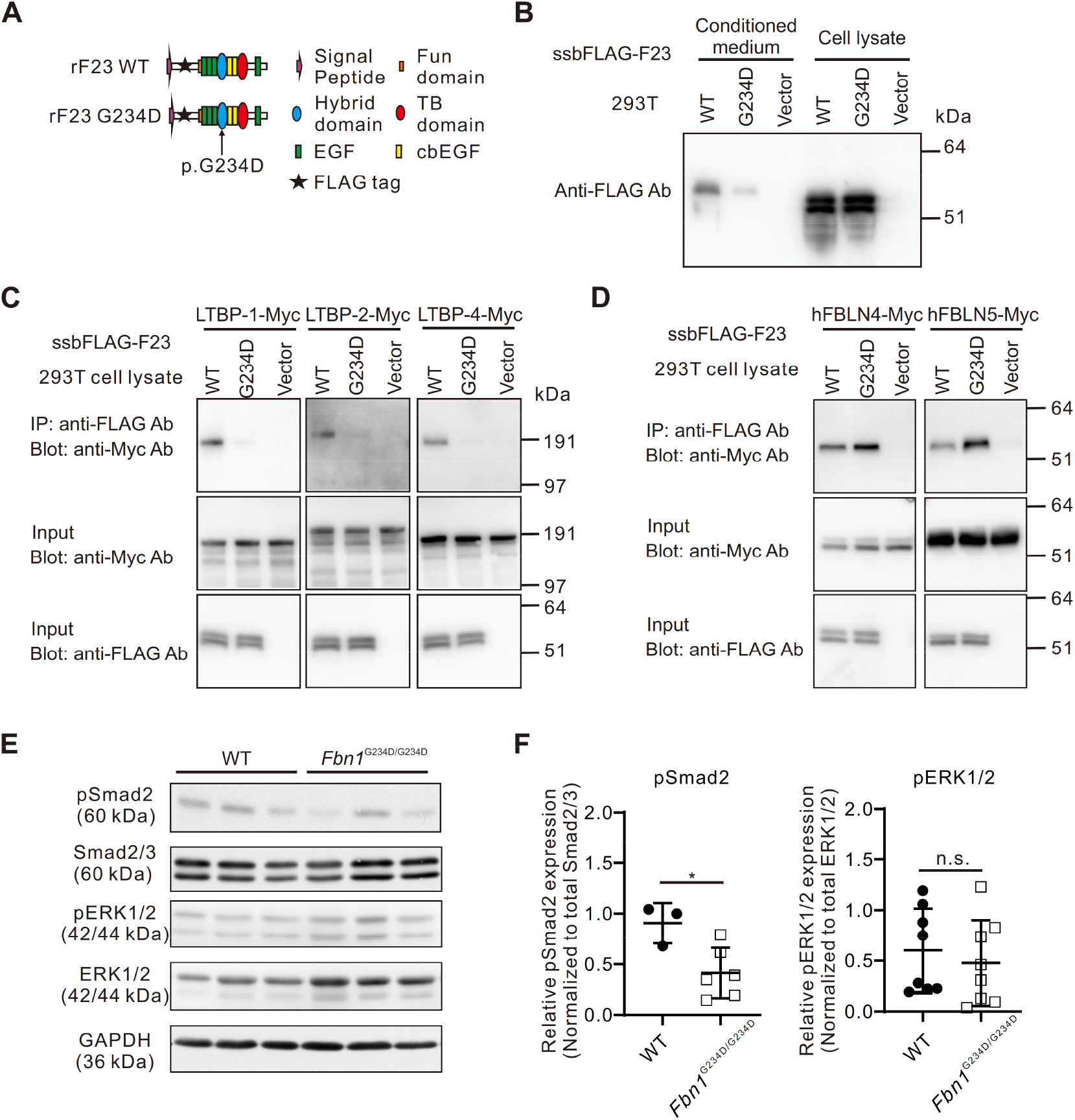
Molecular and biochemical analysis displayed a significant impairment of rF23 G234D in the binding to LTBPs and affected canonical TGFβ pathways. (A) Domain structure of FLAG-tagged partial fibrillin-1 (rF23) and fibrillin-1 mutant (rF23 G234D). (B) Western blot analysis with anti-FLAG antibody using conditioned medium and cells lysates from 293T cells transfected with plasmids encoding the rF23 WT or rF23 G234D. (C) In vitro binding assay using the cell lysates from transfected 293T cells. The lysates were mixed with Myc-tagged LTBPs and immunoprecipitated with anti-FLAG antibodies. (D) In vitro binding assay using the cell lysates from transfected 293T cells. The lysates were mixed with Myc-tagged FBLNs and immunoprecipitated with anti-FLAG antibodies. (E, F) Western blot of total protein extracts from 3-week-old WT and *Fbn1*^G234D/G234D^ ascending aortas. Mean ± SEM, WT: n=3 mice (Smads), n= 8 mice (ERK), *Fbn1*^G234D/G234D^: n=6 mice (Smads), n= 8 mice (ERK), *p<0.05, unpaired *t* test.

It has been proposed that fibrillin-1 regulates the bioavailability of TGFβ in the extracellular microenvironment by binding to LTBP. In the aortic wall of Marfan patients and mice where fibrillin-1 is defective, activation of Smad2/3 and elevated expression of Smad-responsive genes were observed ^11,26^. Our recent study, however, showed removing the first hybrid domain in vivo resulted in microdissection of the aorta and decreased pSmad2 signaling ^7^. Therefore, we examined canonical and noncanonical TGFβ signaling in AD aortas. As Figures 7E and 7F show, no upregulation of pSmad2 or pERK1/2 was observed in AD mice compared with WT mice. Rather, pSmad2 levels in AD aortas were decreased compared to WT aortas. These data indicate that the mutation resulted in failed activation of TGFβ signaling with compromised tethering of LTBPs to Fbn1^G234D^, in contrast to the previously known activation of TGFβ signaling in the Marfan syndrome and related TAA. Our animal model shows a novel mode of regulation of the TGFβ signaling mediated by G234 mutation in the first hybrid domain of fibrillin-1 which contributes new insights into the pathogenesis of aortic dissection in AD mice.

## Discussion

In this study, we established a new mouse model of spontaneous AD by introducing a missense variant of the *FBN1* gene identified in a patient with familial AD. The central pathology of this new mouse model is characterized by spontaneous formation of intimomedial tears in the ascending aorta associated with massive inflammatory cell infiltration. We show that primary defects in fibrillin-1 cause disruption of elastic laminae and dysregulation of subendothelial matrices, which is likely to be responsible for altered responses to flow shear, activation of ECs, and accumulation of monocytes and macrophages, leading to fatal AD. We propose that a major underlying molecular trigger for AD is inflammation, enhanced by the TGFβ-suppressed microenvironment mediated by the G234D mutation in the first hybrid domain of fibrillin-1, which plays a crucial role in initiating intimomedial tears.

### A novel mouse model exhibits preclinical signs of aortic dissection

The intimomedial tears begin to develop after 3 weeks of age in the intimal layer with various severities, involving disruption of a few elastic laminas to multiple medial layers. As long as the external elastic laminae remains intact, a “localized dissection” as described by He et al ^27^ is formed. When the tear develops into a full-thickness one, the surrounding adventitia is strengthened by the accumulation of eosin-positive non-collagenous ECM, resulting in a “contained rupture”. When the adventitia is unable to sustain the aortic wall, it advances to “aortic rupture”. It is of note that multiple intimomedial tears develop simultaneously, and they sometimes coalesce into a large lesion and expand longitudinally and circumferentially, as is demonstrated by the synchrotron-based imaging (Figure 2G, Supplemental Figure 3). We found intimomedial tear volume is significantly higher along the inner curvature in AD mouse. Notably, regional differences in biomechanical properties in the ascending aorta have been reported. In healthy humans, it was shown that failure stresses are smaller longitudinally than circumferentially and more likely to be anteriorly and medially than posteriorly and laterally ^28^. Similar studies using ascending aortic aneurysm samples showed that uniaxial tensile strength was lower at the inner curvature than the posterior wall but orientation of the tissues has more significant impact on material properties, and the aortic wall is more likely to fail in the longitudinal direction ^29^. Interestingly, high wall stresses are observed in Marfan patients with even small aneurysms ^30^. These data indicate heterogeneity of mechanical properties of the aortic wall, which can be influenced by ECM composition and hemodynamics.

### Inflammatory monocytes/macrophages contribute to intimomedial tears

We found that the progression of AD parallels the infiltration of inflammatory cells into the aortic wall. Inflammatory cells, including macrophages, have been observed in the different stages of AD in human patients and older *Fbn1*^mgR/mgR^ mice ^31,32^. Recent single-cell RNA-seq analysis identified both bone-marrow-derived-and residential macrophage populations in dissected lesions ^33,34^. In mouse studies, adventitial macrophages are predominantly observed in AD models that utilize BAPN and AngII with or without hyperlipidemia ^35–37^. A recent study also describes adventitial macrophages in mice carrying the activating mutation of JAK2 V617F in Lyve1+ resident macrophages, which leads to deteriorated AAD with the increased *Il6* and *Tnfa* expression ^38^.

In this study, AD with massive inflammatory infiltrates occurred without any inductions. scRNA-seq and immunofluorescence confirmed accumulation of monocytes/M1-like macrophages in the intima layer of ascending aortas before initiation of AD and these cells are on and underneath ECs, with some of them invading into medial layers. Consistently, we observed upregulation of MMP2/9 activities by in situ gelatin zymography at the site of intimal tears. In addition, upregulation of pNF-kB, which is required for the polarization of pro-inflammatory macrophages and induction of a large number of inflammatory genes ^39^, was observed in the AD aortas as dissection progressed. IL-6 and Stat3 are known to be involved in accelerating macrophage-mediated vascular inflammation, which is suppressed by Socs3 ^36^. In contrast, although CD206+ macrophages increased in the adventitia from 3 weeks of age, these cells did not invade into the media and retained the M2-like marker expression, providing a balance to the pro-inflammatory immune cells recruited to the intimal side.

### Activated endothelial cells serve as an early trigger for aortic dissection

The question remains as to what triggers the accumulation and invasion of monocytes/macrophages into the intimal layer without chemical induction or hyperlipidemic conditions in AD mice. In a physiological context, ECs typically exhibit an elongated and aligned morphology along the direction of blood flow in undisturbed regions. The cell changes seen in AD aortas resemble those seen under turbulent flow, where ECs adopt a cubic shape and display random orientations ^40^. Also, the endothelium in regions prone to atherosclerosis affected by disturbed flow upregulates proinflammatory molecules like MCP1, ICAM-1, and E-selectin, coupled with downregulation of anti-inflammatory transcription factors such as kruppel-like factor (KLF)2 and KLF4 ^41^. It was previously reported that flow-mediated responses were impaired in Marfan patients ^42^ and a negative correlation was observed between flow-mediated dilation and the diameter of ascending thoracic aorta ^43^. In addition, it has previously been shown that Marfan model mice (Fbn1^C1041G/+^) show alteration of EC morphology with a rounder shape and reduced alignment to blood flow ^44^ and reduction of basal NO production in older animals ^45^. The abnormal EC phenotypes with upregulation of ICAM-1 and VCAM-1 are observed as early as 1 week of age before manifestation of the gross phenotype in AD mouse aortas, supporting the notion that the activation of ECs with abnormal mechanosensing contributes to the upregulation of adhesion molecules and triggers the recruitment of bone marrow-derived monocytes/macrophages into the intima.

The biomechanical properties of the ECM environment play an important role in endothelial morphology and functions ^46^. Specifically, increased fibronectin expression has been associated with various vascular diseases, including atherosclerosis ^47^. The upregulation of fibronectin in the subendothelial space represents an early indicator of vascular alterations, potentially contributing to the pathogenesis of atherosclerosis ^48^. Indeed, research using genetically modified mice in which the intracellular domain of integrin α5β1, a fibronectin receptor, is replaced with that of α2β1 collagen receptor, modulates the fibronectin-mediated signaling and attenuates aneurysm formation and improves the survival of *Fbn1*^mgR/mgR^ mice^49^. Interestingly, fibronectin was also elevated in subendothelial lesions and medial layers in AD mice and upregulation of laminin and collagen IV was detected, indicating the alteration of ECM composition due to an alteration of fibrillin-1 microfibrils in the subendothelial matrix. It will be informative to identify the cell-type responsible for producing ectopic ECM.

### The regulation of the TGFβ signaling pathway by FBN1

Numerous missense variants have been identified in the *FBN1* gene and the intriguing observation of various phenotypes resulting from these variants raises questions about their connection to the regulation of the TGFβ signaling pathway (Reviewed in ^10^). Using an in vitro system, we show that the fibrillin-1^G234D^ protein lacks the capacity to bind LTBP-1, -2 and -4. The hybrid domain where G234 is located contains the LTBP-1 binding site and disruption of fibrillin-1 has been suggested to increase the bioavailability of TGFβ, thus increasing the TGFβ-mediated signaling in Marfan syndrome ^11,50^. However, the early in vivo study deleting the Hybrid domain 1 (*Fbn1*^H1Δ/+^ or *Fbn1*^H1Δ/H1Δ^) showed normal assembly of microfibrils without aortic aneurysms ^51^. A recent finding however revealed that *Fbn1*^H1Δ/+^ mice develop microdissections with mast cells present in the aortic wall, associated with a decreased expression of TGFβ downstream molecules ^7^. Similarly in AD aortas, the mutant fibrillin-1 with the inability to bind LTBPs and significantly decreased pSmad2 levels provides the permissive microenvironment for inflammatory cell infiltration. Indeed, destruction of the aortic wall and the inflammatory cell-dominant phenotype in AD aortas resembles that of SMC-specific TGFβ receptor type II deletion in adult mice, except that macrophage infiltration is mainly observed in the media in this case ^52^. Additionally, a recent ultrastructural analysis of native fibrillin microfibrils has revealed that the loss of the first hybrid domain in FBN1 leads to the formation of abnormal microfibril structures ^53^, which has a strong impact on elastic fiber assembly. Taken together, it is reasonable to speculate that the G234D mutation disrupts the formation of microfibrils and impacts elastic lamella stability as well as downregulates TGFβ signaling. Further study of the hybrid domain’s involvement in modulating this pathway will unveil the mechanisms underpinning the diverse outcomes observed in various *Fbn1* mutants.

### Implication to human aortic dissection

As for molecular pathology of the *FBN1* p.Gly234Asp variant, we found that the *FBN1* G234V variant was also reported in a large Danish Marfan family with high penetrance of fatal AD ^54^. Although precise clinical information was not available in this family and the pathogenicity of the G234D and the G234V variant according to the ACMG/AMP standards ^55^ need proper curation, our current data and G234V mutation emphasize the importance of the glycine residue at this location. Indeed, it is reported that the G-F/Y dipeptides are involved in interdomain interactions ^56^, and changing glycine to aspartic acid or valine is likely to cause conformational changes that are detrimental to the function of fibrillin-1.

Recent transcriptomic studies comparing the healthy aorta and dissected aortas in sporadic TAAD and genetic aortic disease have pointed to the contribution of inflammatory cells ^57^. Preclinical studies targeting the inflammation pathway have been reported for AA and AAD using a wide range of strategies, including indomethacin to inhibit COX-1 and COX-2 ^37^, neutralizing antibodies against GM-CSF ^58^ and an M-CSF receptor tyrosine kinase inhibitor Ki20227 to inhibit macrophage accumulation ^33^ as well as IL-11 receptor antagonist X209 ^59^. Our spontaneous AD mouse model indicates bone-marrow-derived monocytes/macrophages as a critical component of AD pathology and opens an avenue to develop a focused strategy directly targeting these inflammatory cells. Although the AD mice rapidly develop AD at a young age, this model underscores the multifaceted nature of AD pathology mimicking some aspects of the human condition and serves as a useful model to validate potential drug targets for AD, such as protein kinase C-beta inhibitor recently proposed for vascular Ehlers-Danlos syndrome ^8^. Taken together, our new mouse model can provide a foundation for further exploration of the molecular mechanisms involving inflammation during the initiation of AD in familial or sporadic aortic aneurysms and dissection and for the development of specifically targeted therapies.

## Nonstandard Abbreviations and Acronyms

AD: aortic dissection
TAA: thoracic aortic aneurysm
TAAD: thoracic aortic aneurysm and dissection
AA: aortic aneurysm
FBN1: fibrillin1
TGFβ: transforming growth factor-beta
LTBP: latent TGFβ binding protein
ECM: extracellular matrix
EC: endothelial cell
SMC: smooth muscle cell

## Acknowledgment

We thank the Laboratory Animal Resource Center at the University of Tsukuba for their excellent animal care. We also thank Yoan Cherasse for a custom-built computer for sc-RNA-seq analysis, Keisuke Shirokura for teaching us En Face preparation and immunostaining of the ascending aorta, Dieter Reinhardt for *FBN1* expression plasmid, Bart Lambrecht, Haruka Ozaki, Kaori Sugiyama and Yoshito Yamashiro for discussion. This study was supported in part by the NOVARTIS Foundation (Japan) for the Promotion of Science (to K.K.), SENSHIN Medical Research Foundation, Japan Agency for Medical Research and Development (grant number 23ek0109553h003 and 23ek0210183h0001), Grant-in-Aid for Scientific Research (A, grant number 23H00431), and Grant-in-Aid for Fund for the Promotion of Joint International Research (Fostering Joint International Research(B)) (grant number 21KK0151) (all to H.Y.).

## Author contributions

K.K. and E.M. performed mouse experiments, immunohistochemistry, cell culture, imaging, and data analysis. S.K. identified a new fibrillin-1 variant and provided patient data. K.A, M.A.A.S., and R.I. performed scRNA-seq and analyzed the data. M.T.R.D.C.C. performed in situ zymography experiment and analyzed the data. E.R. performed mouse phenotypic analysis and analyzed the data. M.T. and T.N. performed biochemistry experiments and analyzed the data. K.M. and S.M. generated experimental animals. M.A.A.S., V.D., and P.S. performed synchrotron experiments and analyzed the data. L.M.M, J.D.B., and L.Y.S. provided expertise on aortic diseases and fibrillin-1 biology and interpreted the results. K.K. and H.Y. designed experiments, interpreted the results, supervised the project, obtained funding, and wrote the manuscript. All authors edited the manuscript.

